# Accurate nucleic acid-binding residue identification based on domain-adaptive protein language model and explainable geometric deep learning

**DOI:** 10.1101/2024.12.11.628078

**Authors:** Wenwu Zeng, Liangrui Pan, Boya Ji, Liwen Xu, Shaoliang Peng

## Abstract

Protein-nucleic acid interactions play a fundamental and critical role in a wide range of life activities. Accurate identification of nucleic acid-binding residues helps to understand the intrinsic mechanisms of the interactions. However, the accuracy and interpretability of existing computational methods for recognizing nucleic acid-binding residues need to be further improved. Here, we propose a novel method called GeSite based the domain adaptive protein language model and explainable E(3)-equivariant graph convolution neural network. Prediction results across multiple benchmark test sets demonstrate that GeSite is superior or comparable to state-of-the-art prediction methods. The performance comparison on low structure similarity and newly released test proteins demonstrates the robustness and generalization of the method. Detailed experimental results suggest that the advanced performance of GeSite lies in the well-designed nucleic acid-binding protein adaptive language model. Meanwhile, interpretability analysis exposes the perception of the prediction model on various remote and close functional domains, which is the source of its discernment. The data and source code of GeSite are freely accessible at https://github.com/pengsl-lab/GeSite.

## INTRODUCTION

Protein-nucleic acid interactions serve as a fundamental role in various biological processes like gene regulation and expression in organisms (1). Understanding these interactions is crucial for studying protein function and facilitating drug development. Accurate identification of nucleic acid-binding residue (NBS) is a key step in elucidating the underlying mechanisms of these interactions. Traditional biological wet-lab experimental methods, including chromatin immunoprecipitation on microarrays, nuclear magnetic resonance, and X-ray crystallography, have significantly advanced the study of protein-nucleic acid interactions. However, these methods are limited by high costs and long lead times, making it impractical to determine all NBSs across the vast number of protein sequences in the post-genomic era. As of 14 November 2024, there are 833,496,420 protein sequences recorded in the UniParc database (2), while the number of resolved protein-DNA (or RNA) complex structures recorded in the PDB Database (3) is only 226,193 (or 225,010). Although recent methods like AlphaFold3 (4) and RoseTTAFold All-Atom (5) attempted to predict the 3D structure of protein-nucleic acid complexes directly, the accuracy remains suboptimal due to the limited number of known complex structures that such computational methods heavily depend on. Accurate binding sites can further assist in modeling protein-nucleic acid complexes. Consequently, the development of rapid and accurate method for predicting NBS remains essential. Numerous computational methods have been proposed to address this challenge. In the early days, researchers used statistical and machine learning-based methods to capture conserved information of NBS for prediction. Despite the progress made, these methods generally suffered from poor accuracy and generalizability. In the past decade, deep learning-based methods have gained significant attention. These methods capture the intricate non-linear relationships between the sequence, structure, and function of proteins, enabling highly accurate NBS prediction. Depending on the feature sources employed, these methods can be broadly classified into two categories: sequence-driven methods, e.g., iDRNA-ITF (6), ESM-NBR (7), ULDNA (8), CLAPE (9), hybridRNAbind (10), and HybridDBRpred (11); as well as structure-driven methods, e.g., GraphSite (12), GraphBind (13), CrossBind (14), and EquiPNAS (15). Sequence-driven methods typically explore evolutionary information through primary sequences to identify nucleic acid-binding residues. While these methods are convenient and easily scalable, their accuracy is often limited due to the inherent difficulty of extracting useful discriminative information directly from primary sequences. On the other hand, structure-driven methods leverage the high conservatism and specificity of NBS on 3D structure, potentially achieving better performance. However, the longstanding lack of high-quality protein 3D structures has hindered the scalability of such methods. Promising to change this dilemma are recent major advances (16-18) in deep learning-based protein 3D structure prediction like AlphaFold2 (19). Using the predicted 3D structure as a substitute can complement the inadequacy of the resolved native structure. Moreover, the role of protein language model (PLM) in studying protein function and structure (20,21) provides new insights into the analysis of protein-nucleic acid interactions. Integrating protein structure features with knowledge from PLM is expected to significantly advance the study of protein-nucleic acid interactions.

In recent years, several methods have been developed to predict nucleic acid-binding residues based on protein structure (whether native or predicted) and universal PLM. For example, CrossBind (14) predicts NBS by integrating amino acid-level PLM and atom-level feature representation. In GraphBind (13), a local graph centered on the target residue is first constructed to represent the structural context; then a hierarchical graph neural network is used to learn embedded rules to recognize NBS. Liu *et al*. proposed a DNA-binding residue predictor called CLAPE that combines the PLM ProtTrans (22) and the contrastive learning strategy. In ULDNA (8), Zhu *et al*. first extracted feature representations from two single-sequence PLM, i.e., ESM2 and ProtTrans, and one multiple sequence alignment (MSA) transformer model, i.e., ESM-MSA (23), then fed into an LSTM-attention network for prediction. In EquiPNAS (15), ESM2 feature embedding, MSA representation extracted from AlphaFold2, and a variety of local structural features serve as inputs to train an E(3)-equivariant graph neural network (EGNN) (24).

Despite the good results achieved by the aforementioned methods, there is still room for improvement. First, PLM-based NBS prediction methods typically extract sequence embedding as feature representations directly from the original universal PLM trained on massive general protein sequences. While these PLMs are excellent for characterizing properties common to all protein families, such as tertiary structure, they are under-explored for specificity when focusing on particular families like Nuclear Receptor and Forkhead, which are typical DNA-binding protein (DBP) families. The ability of proteins to bind to nucleic acids mainly comes from a structural domain folded by a small, highly conserved sequence motif. Further probing of these typical nucleic acids-binding motifs will undoubtedly further enhance the ability of PLM to characterize nucleic acids-binding protein (NBP), thereby improving NBS prediction accuracy. Second, to capture contextual information about spatial structures, previous methods (12,13) use several GNN variants such as graph transformer and Gate Recurrent Unit (GRU)-based GNN (25), which, despite their good performance, do not provide visualization and interpretability, making it difficult to intuitively understand what the model has actually learned.

In this study, we propose a novel structure-based NBS prediction method named GeSite based on DNA- and RNA-binding protein domain-adaptive PLM and explainable EGNN. In GeSite, we directly utilized our previous study, ESM-DBP (26), as the DBP adaptive PLM to extract sequence embedding as input feature for DNA-binding residue (DBS) prediction. For RNA-binding protein (RBP) adaptive PLM, similar to ESM-DBP, we collected 459,656 non-redundant RBP sequences to fine-tuning the parameters of the last five transformer blocks of ESM2, resulting in ESM-RBP for RNA-binding residue (RBS) prediction. Subsequently, the MSA file is generated using the HHblits (27) tool to search the Uniclust30 database (28) and fed into the ESM-MSA model to obtain the embedding matrix as the representation of evolutionary information of protein sequence. The embedding matrix extracted from the well-trained ESM-D/RBP is concatenated with the output of ESM-MSA to serve as the feature representation of each protein sequence in the benchmark dataset. Finally, a protein graph based on residue contact map is constructed for input into EGNN to predict the binding probability of each residue. The prediction results on three widely used benchmark test sets show that the performance of GeSite on both the DBS and RBS are better than or comparable to state-of-the-art (SOTA) methods. The AUC values of GeSite on DNA-129_Test, DNA-181_Test, and RNA-117_Test are 0.941, 0.919, and 0.861, which are 0.75, 0.22, and 9.96% higher than that of the second-best method separately. The performance comparison with two SOTA structure-based methods on the low structure identity test proteins and a newly collected RBP test set ulteriorly demonstrates the generalization and robustness of the proposed method. The experimental results demonstrate that the superior performance of the proposed method is based on the domain-adaptive PLM that provides better sequence characterization of NBS than the universal PLM. In addition, interpretability analysis and visualization of the prediction model reveal the sensitivity of GeSite to various nucleic acids-binding domains, which is the source of its recognition ability. Especially, the focus on remote functional domain is also observed on the DNA ligase of African swine fever virus.

## METHODS

### Benchmark datasets

As in the ESM-DBP, to train an RNA-binding protein domain-adaptive PLM, we first collected 9,743,473 redundant RBPs (up to February 6, 2024) from UniProtKB database (29); then, to prevent the model from overfitting the RBP family with high redundancy, CD-HIT tool (30) was used to remove those high similarity sequence using a cluster threshold of 0.4 and remaining 459,656 non-redundant RBPs named UniRBP40 as the pretraining data set.

To facilitate the verification of the performance of GeSite, two nucleic acids-protein binding datasets used extensively in previous studies are employed in this study. Specifically, for DNA-binding residues prediction, one training data set named DNA-573_Train and two independent test sets, i.e., DNA-129_Test and DNA-181_Test, are employed; for RNA-binding residues prediction, one training data set RNA-495_Train and one test set RNA-117_Test are employed. Of these benchmark datasets, DNA-573_Train, DNA-129_Test, RNA-495_Train, and RNA-117_Test are collected from BioLip database (31,32) by Xia *et al*. in GraphBind (13); DNA-181_Test is the newly released protein (from 6 December 2018 to August 2021) in BioLip database and collected by Yuan *et al*. in GraphSite (12). To ensure a fair and objective performance evaluation, the CD-HIT program with a cluster threshold of 0.3 was used to remove protein sequences that hold high sequence identity between the test sets and training sets.

In addition, to more fully evaluate the generalizability of the present method, we constructed a new RBS test set named RNA-285_Test from newly released protein-RNA complexes. Specifically, we first collected all 160,800 RNA-binding entries including 6,041 RBPs (128,464 chains) from the BioLip database from 6 December 2018 to 19 October 2024 (independently of RNA-495_Train and RNA-117_Test); second, CD-HIT tool (cluster threshold 0.4) was utilized to cluster all chains and the representation sequences are selected; third, those sequences with high identity to RNA-495_Train and RNA-117_Test were removed using CD-HIT-2D tool with the threshold of 0.4; finally, to remove redundancy on the structure, we used the US-align tool (33) to calculate the maximum TM-score for each remaining chain against the existing RBS dataset (normalization by longer chains), and those chains with a maximum TM-score greater than 0.5 were filtered out, remaining 285 RNA-binding chains ultimately. The detailed components of these datasets are listed in Table 1.

**Table 1.** Composition of the training and testing data sets.

| Type | Dataset | $N_{\text{protein}}^a$ | $N_{\text{posi}}^b$ | $N_{\text{nega}}^c$ | $\text{Ratio}^d$ |
| --- | --- | --- | --- | --- | --- |
| DBS | DNA-573_Train | 573 | 14,479 | 145,404 | 0.100 |
|  | DNA-129_Test | 129 | 2,240 | 35,275 | 0.064 |
|  | DNA-181_Test | 181 | 3,208 | 72,050 | 0.044 |
| RBS | RNA-495_Train | 495 | 14,609 | 122,290 | 0.119 |
|  | RNA-117_Test | 117 | 2,031 | 35,314 | 0.058 |
|  | RNA-285_Test | 285 | 4245 | 41399 | 0.102 |

a. Number of proteins.
b. Number of binding residues.
c. Number of non-binding residues.
d. $\text{Ratio} = N_{\text{posi}}/N_{\text{nega}}$ .

### Domain-adaptive protein language model

Recent advances in protein function and structure prediction based on PLM demonstrate that PLM learns amino acid dependencies to efficiently characterize protein sequence. This learning paradigm is largely inspired by the BERT-based large language model (LLM) in the field of natural language processing (NLP) (34). A recent study (35) about NLP showed that domain-adaptive pretraining can provide significant gains in downstream task performance. This idea can be naturally transferred to PLM. In ESM-1b (21), Rives *et al*. mentioned that the PLM after self-supervision learning encodes the MSA knowledge into the sequence representation. It is easy to imagine that if PLM is ulteriorly trained on particular protein families, the sequence characterization of these particular families will be further improved. In our previous study ESM-DBP, through domain-adaptive pretraining on massive DBP sequence data, the proposed DBP domain-adaptive PLM improves prediction performance and outperforms SOTA methods on several DBP-related tasks. Here, similar to ESM-DBP, to construct ESM-RBP, we continue to train the ESM2 model consisting of 33 transformer blocks with 650 million parameters by randomly masking and then predicting 15% residues of each sequence in UniRBP40 (see Figure 1A). Slightly different, since the number of protein sequences in UniRBP40 (459,656) is greater than UniDBP40 (170,264), we increase the number of updatable transformer blocks to 5 about 100 million parameters. The first 28 transformer blocks hold the fundamental biological knowledge that ESM2 learned from about 65 million general sequences in UniRef50, and the last 5 transformer blocks possess the RBP-specific knowledge learned through continued training from massive RBPs. In the pretraining phase, the cross-entropy and Adam optimizer are used to calculate the loss and update the parameters, respectively. Each input sequence consists of 512 tokens, short sequences are filled with token of <pad>, and sequences longer than 512 are split. The batch size is set to 230 according to the available memory. The model is optimized for about 35K steps and took about 3 days on two Tesla A40 GPUs with 48G memory.

**Figure 1.**
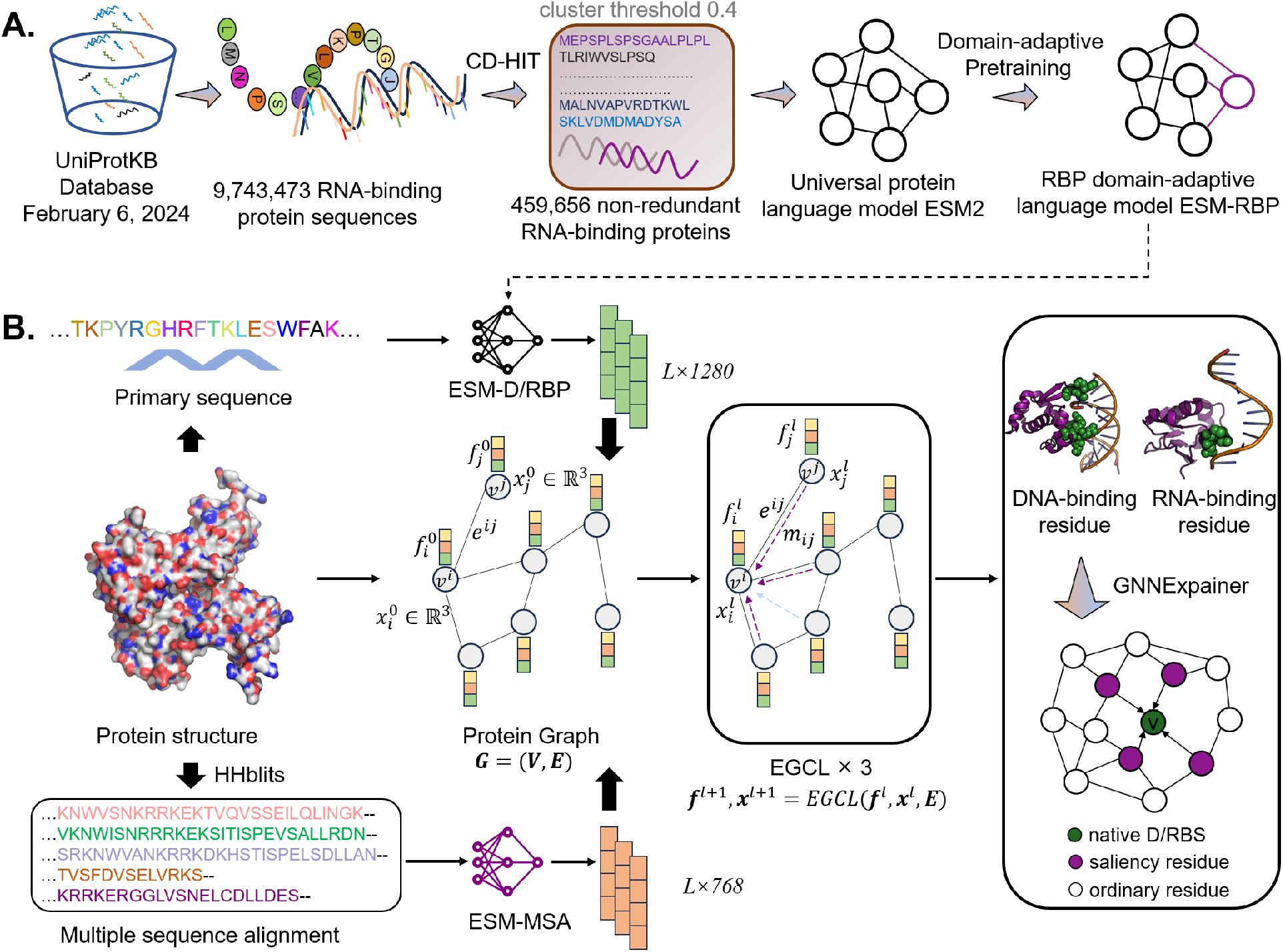
The overall architecture of GeSite. **(A).** flow chart of construction of domain-adaptive protein language model; **(B)**. nucleic acid-binding residue prediction based on ESM-D/RBP and E(3) equivariant graph neural network. ESM-DBP (26) is our previous study, and ESM-RBP is trained de novo in this study.

### Protein Graph Representation

We converted the protein structure into a graph representation to learn spatial feature of target residue for NBS prediction. Specifically, a protein of length *s* is represented by a graph ***G =* (*V***,***E*)**, where ***V =***{ν^0^,…, ν^*i*^,…, ν^*s*−1^} represents the set of all residue nodes; *e*^*ij*^ ∈ ***E*** represents the set of edges of interacting residues. Detailed descriptions of generation of node and edge features are as follows:

#### Node feature

We treat each residue as a node in the graph ***G***. For each protein sequence of length *ss*, we first input it into the ESM-D/RBP to generate an embedding matrix of size *s*× 1280; then the Multiple Sequence Alignment (MSA) file is generated by searching Uniclust30 database (28) using HHblits tool (27) and fed to ESM-MSA (23) to obtain an embedding matrix of size *s*× 768 for portraying the evolutionary information of a protein (see Supplementary Text S2 in detailed); finally, these two embedding matrixes are concatenated into the feature representation matrix ***f =***{*f*_0_,…, *f*_*i*_, …, *f*_*s*−1_} of size *s*× 2,048 of the target protein. Each feature of residue node ν^*i*^ is a vector *f*_*i*_ of length 2,048.

#### Edge feature

Edges portray associations between nodes and are an important source of spatial information about neighboring residues. Here, we define edge for residue pairs that are close in spatial distance. In particular, if the Euclidean distance between the C*a* atoms of two residues is less than 14Å, then they are considered to be in contact. The edge feature *e*^*ij*^ of target residue pair (*R*^*i*^, *R*^*j*^) is |*R*^*i*^, *R*^*j*^|/*D*_*max*_, where |*R*^*i*^, *R*^*j*^| means Euclidean distance between the C*a* atoms of residues *R*^*i*^ and *R*^*j*^; *D*_*max*_ means the maximum residue distance in the target protein.

In addition to node and edge features, EGNN introduces coordinate feature *x*_*i*_ for each node *v*^*i*^. Here, *x*_*i*_ is denoted by the three-dimensional Cartesian coordinates of C*a* atom of residue *v*^*i*^. EGNN retains equivariance to rotations and translations on coordinate set ***x*** and to permutations on node set ***V*** (24).

### Implement of E(3) Equivariant Graph Neural Network

GNN and the variants are widely used to capture knowledge of protein 3D structure since they are adept at extracting spatial contextual embeddings of neighboring residues. The equivariant properties of EGNN in rotations, translations, reflections and alignments allow it to maintain the symmetry of the graph structure. In this study, EGNN is employed as prediction model to learn more abundant structure representation of protein than traditional graph convolutional network (GCN). See Figure 1B, EGNN is composed of three Equivariant Graph Convolutional Layers (*EGCL*) which takes the residue node feature set ***f***^*l*^, edge set ***E***, and coordinate set ***x***^*l*^ as input and performs a transformation *EGCL***(*f***^*l*^, ***x***^*l*^, ***E***) as follows:

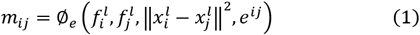

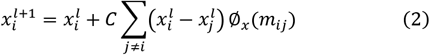

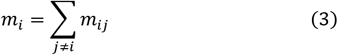

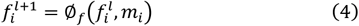

where ∅_*e*_ and ∅_*f*_ mean edge and node operations respectively based on Multilayer Perceptron (MLP) and *Swish*() activation function which are similar to typical GCN; *C* ***=***1/(*s*− 1) means taking the average; ∅_*x*_= {*Linear*() → *Switch*() → *Linear*()} converts *m*_*i*_ into a scalar value as the weight of relative difference 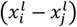. The outputted node embedding ***f***^*l*+1^ and coordinate set ***x***^*l*+1^ are used as inputs of the next *EGCL*(). The involvement of the coordinate information in the updating of the node embedding is the main difference that distinguishes EGNN from traditional GCN, and is the source of its equivariance on rotations and translations.

The GeSite model is implemented using the Pytorch and the DGL frameworks (36). Limited by available memory size, the batch size is set to 1, that is, each batch uses one protein graph for forward and back propagation. The cross-entropy and AdamW optimizer (37) with a learning rate of 1e-4 are used to portray the loss and optimize parameters. Considering the category imbalance, the loss weights for the positive and negative samples are 0.7 and 0.3, respectively. To avoid overfitting, regularization with a coefficient of 1e-04 is employed to restrict the parameters. The entire training process lasted 50 epochs on a Tesla V100 GPU with the memory of 16G. Notably, our GeSite consists of two independent single-task models predicting DNA- and RNA-binding residues, respectively.

### Architecture of GeSite

As shown in Figure 1, the architecture of the proposed GeSite can be roughly divided into three steps:

*Step 1. Domain-adaptive pretraining*: similar to the previous study ESM-DBP, the RBP domain-adaptive PLM i.e., ESM-RBP, is well-trained based on universal PLM ESM2 using the pretraining dataset of UniRBP40 contained 459,656 non-redundant RNA-binding sequences;

*Step 2. Constructing the protein graph*: the protein 3D structure is mapped into a protein graph with residues as nodes and feature embeddings extracted from ESM-D/RBP and ESM-MSA, respectively, as node features. Residue pairs with Euclidean distances less than 14Å are given edges with normalized residue distances as weights.

*Step 3. Predicting NBS based EGNN*: the protein graph is fed into an EGNN model consisting of three EGCL to predict the NBS. The output of each node of the last EGCL is used as the probability that the target residue is predicted as an NBS.

## RESULTS

### Role of Domain-adaptive Language Model

To demonstrate the advantages of the NBP domain-adaptive language models, i.e., ESM-DBP and ESM-RBP, over ESM2 for the task of NBS prediction, we replace the ESM-D/RBP sequence embeddings in GeSite with the original ESM2 feature embeddings and then retrain the GeSite model for comparison. From Table 2, the MCC values of GeSite using ESM-D/RBP feature embeddings on DNA-129_Test, DNA-181_Test, and RNA-117_Test are 0.522, 0.389, and 0.326, which are 8.75, 7.45, and 14.38% higher than those of GeSite using ESM2 feature embeddings respectively. Considering other evaluation indexes, the model using ESM-D/RBP is also better than that using original ESM2 at prediction performance. Taking prediction results on DNA-129_Test as an example, the Spe, Rec, Pre, F_1_, AUC, and AP values of the former are 0.948, 0.618, 0.431, 0.508, 8.07, 1.51, and 8.68% than those of the latter respectively.

**Table 2.** Performance comparison of the sequence representation of original ESM2 and ESM-D/RBP on independent test sets.

| Test set | Feature | Spe | Rec | Pre | F<sub>1</sub> | MCC | AUC | AP |
| --- | --- | --- | --- | --- | --- | --- | --- | --- |
| DNA- | ESM2<sup>a</sup> | 0.948 | 0.618 | 0.431 | 0.508 | 0.480 | 0.927 | 0.518 |
| 129_Test | ESM-DBP<sup>b</sup> | **0.956<sup>c</sup>** | **0.638** | **0.481** | **0.549** | **0.522** | **0.941** | **0.563** |
| DNA- | ESM2 | **0.931** | 0.581 | 0.274 | 0.373 | 0.362 | 0.904 | 0.334 |
| 181_Test | ESM-DBP | 0.925 | **0.650** | **0.279** | **0.391** | **0.389** | **0.919** | **0.367** |
| RNA- | ESM2 | 0.838 | **0.652** | 0.188 | 0.293 | 0.285 | 0.837 | 0.245 |
| 117_Test | ESM-RBP<sup>b</sup> | **0.909** | 0.550 | **0.258** | **0.352** | **0.326** | **0.861** | **0.271** |

a. The prediction results are obtained by replacing the ESM-D/RBP embedding in GeSite with the original ESM2 embedding.
b. For the DBS and RBS predictions, we use DBP and RBP domain-adaptive PLMs to extract sequence embeddings, respectively.
c. The best-performing results are highlighted in bold.

Apart from that, for a more intuitionistic comparison at the protein level, Figure 2 illustrates a head-to-head comparison of the proteins in the three test sets. By looking at Figure 2, regardless of the test set, on most of the proteins, GeSite has better prediction results than the ESM2. For example, for DBS predictions, 77 of the 129 DBPs in DNA-129_Test have higher MCC values for GeSite than ESM2; for RBS prediction, 69 of the 117 RBPs in RNA-117_Test have higher MCC values for GeSite than ESM2. We also note that most of the RBPs (Figure 2C) are located closer to the bottom left than the DBPs (Figures 2A and 2B), implying that the overall predictive performance of the RBS is lower than that of the DBS. There are two main potential reasons for this: first, the number of DBPs (573) in the training set is higher than that of RBPs (495); and second, the binding patterns of proteins and RNAs are more complex and are difficult to be captured well by the model. Nevertheless, in Figure 2C, it is still intuitively clear that GeSite using the ESM-RBP sequence embedding performs much better than that using the original ESM2 embedding representation.

**Figure 2.**
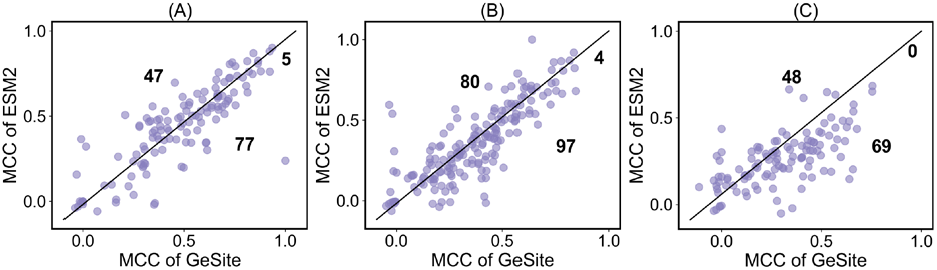
Head-to-head comparison of the MCC values of GeSite and ESM2 on the three test sets at the protein-level. Each dot represents a protein. The number in the upper triangle (or lower triangle) indicates the number of samples located in this region. Points on the diagonal indicate that the two methods share equal MCC values for that protein. **(A)**. on DNA-129_Test; **(B)**. on DNA-181_Test; **(C)**. on RNA-117_Test.

The above experimental results show that the domain-adaptive PLM pays attention to more in-depth identification knowledge of nucleic acid-binding patterns which provides a better sequence characterization of NBP after the domain-adaptive pretraining on a large number of nucleic acid-binding sequences, and thus improves the NBS prediction performance.

### Performance of different network architectures

We utilized different neural network algorithms to train the proposed GeSite model to highlight the suitability of EGNN in the two prediction tasks of this study. Concretely, five structure-based GNNs, i.e., Graph convolutional neural network (GCN), Graph Sample and Aggregate (GraphSAGE) (38), Geometric Vector Perceptrons Graph neural network (GVP-GNN) (39), Graph Attention Network (GAT) (40), and EGNN, as well as three sequence-based networks, i.e., BiGRU, BiLSTM, and self-Attention (41) are employed to perform NBS prediction tasks on the DNA-573_Train and RNA-495_Train over the 10-fold cross validation, respectively. Except for the core algorithm, which is different, the training strategies of these models remain consistent with standard GeSite. Figures 3A and 3B show the PRCs (left part) and ROCs (middle) of these eight models on two data sets separately. The ROC embodies a comprehensive assessment of the predictive performance of NBS and non-NBS. The PRC, on the other hand, is more concerned with sensitivity to positive samples. Visually, for the ROC metric, EGNN achieves the highest score regardless of the prediction task. The AUC value of EGNN on DNA-573_Train (or RNA-495_Train) is 0.918 (or 0.881), which is 1.10, 1.32, 10.74, 13.19, 1.21, 0.66, and 4.68% (or 1.50, 2.09, 12.23, 13.09, 1.97, 3.65, and 13.53%) higher than that of GraphSAGE, GVP-GNN, GCN, GAT, BiLSTM, BiGRU, and Self-attention respectively. This indicates that the overall comprehensive performance of the EGNN model outperforms other algorithms, at least in this study. Observing the PRC curves, the EGNN model obtains the highest and second-best AP values on the RBS (0.517) and DBS (0.560) data sets respectively, which demonstrates the sensitivity of EGNN to the NBS. Although EGNN has a slightly lower AP score on the DBS prediction than BiGRU, it is nevertheless 10.23% higher on the RBS prediction. Consequently, EGNN is considered a more suitable option. Additionally, we also visualized the balance metric MCC score (refer to Figure 3C). Similarly, EGNN achieves the highest MCC scores of 0.514 and 0.454 on both tasks emphasizing the superiority of EGNN again. Comparing structure-based and sequence-based methods, no significant superiority of the former over the latter is observed. After all, protein structures are heavily determined by sequences, and our adaptive language model is able to refine valid knowledge from primary sequences for NBS prediction. Among five structure models, GCN and GAT perform relatively poorly; among three sequence models, Self-attention is worse than BiLSTM and BiGRU. For GCN, the potential reason may be that its ability to extract discriminatory knowledge from the local structure of proteins to predict NBS is somewhat lacking. For algorithms such as Self-attention and GAT, which are suitable for scenarios with a large amount of data, we hypothesize that their mediocre prediction results are caused by insufficient training data. We also note that, as an equivariant GNN like EGNN, GVP-GNN performs slightly lower than EGNN. The remaining indexes see Supplementary Table S1.

**Figure 3.**
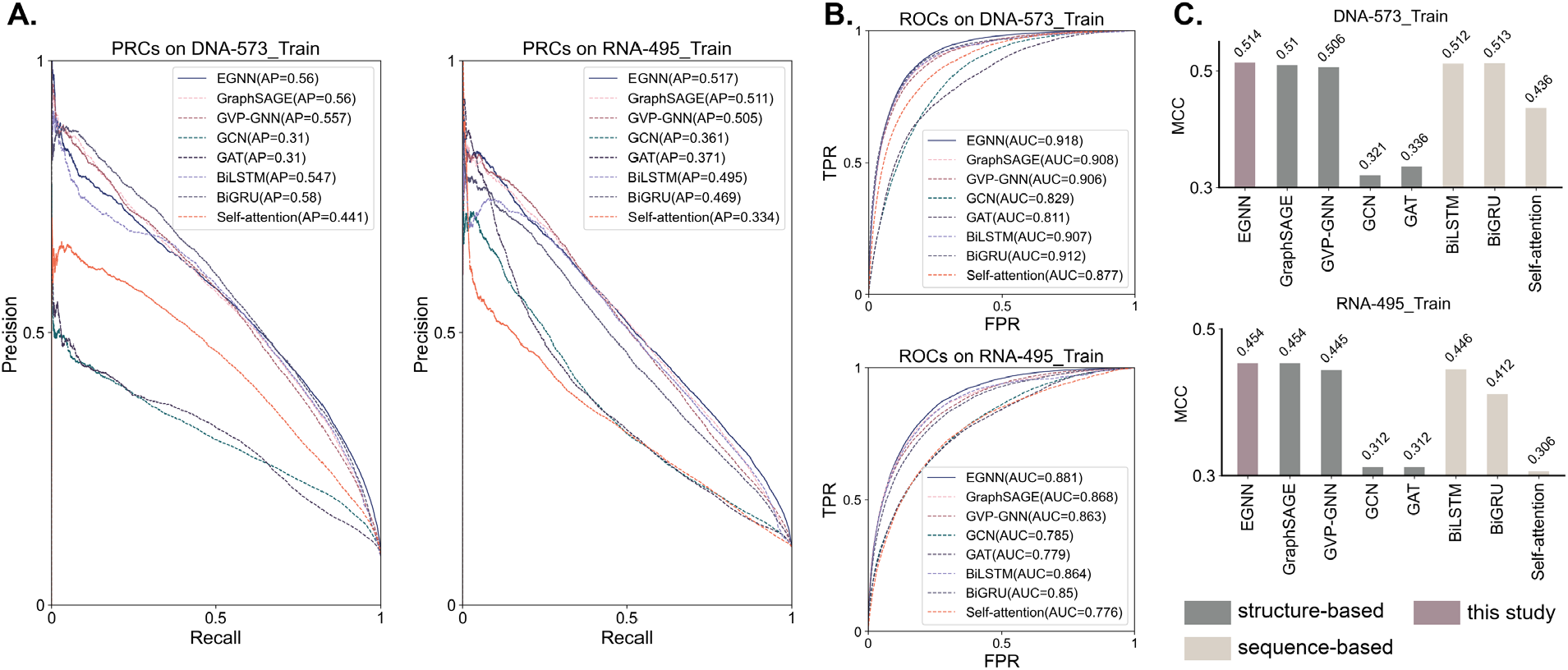
Performance comparison of five structure-based and three sequence-based neural network algorithms on training data sets DNA-573_Train and RNA-495_Train over 10-fold cross validation. **(A)**. PRC; **(B)**. ROC; **(C)**. MCC.

In summary, the EGNN model learns more discriminatory information to distinguish between NBS and non-NBS than other networks in this study. From the experiment results, EGNN better perceives the structural nucleic acid-binding specificity knowledge compared to other GNNs. Compared to sequence-based models, EGNN is also superior in accuracy. Consequently, EGNN is utilized as the core prediction algorithm of GeSite for NBS prediction.

### Comparison with state-of-the-art methods

To future demonstrate the validity of the proposed GeSite for predicting NBS, 9 DBS predictors (EquiPNAS, GraphBind, GraphSite, ESM-NBR, CLAPE, DRNAPred (42), ULDNA, iDRNA-ITF, DNApred (43), and hybridDBRpred) and 7 RBS predictors (EquiPNAS, CLAPE, iDRNA-ITF, GraphBind, ESM-NBR, DRNAPred, Pprint2 (44), and hybridRNAbind) are employed as control methods.

Table 3 shows the detailed prediction results of these methods on three independent test sets, i.e., DNA-129_Test, DNA-181_Test, and RNA-117_Test. From Table 3, in both DBS and RBS prediction, GeSite demonstrates outstanding prediction performance that is superior to most of the other methods. Specifically, for DBS prediction, taking the DNA-129_Test as an example, the MCC value of GeSite is 0.522, which is 7.85, 0.58, 14.47, 34.19, 236.77, 19.72, 34.88, 51.3, and 198.29% higher than those of other methods, respectively. For RBS prediction, the MCC value of GeSite on RNA-117_Test is 0.326, achieving improvements of 35.83, 38.14, 94.05, 76.22, 1317.39, 77.17, and 171.66% over the control methods separately. The leading performance in other indicators also demonstrates the comprehensive performance of the proposed method. To observe the difference between the proposed GeSite and the existing methods, the PCC and *p*-value are calculated. Specifically, we calculate PCC using the probabilities that all residues in the test set are predicted to be NBSs. While for *p*-value, the probability of native NBS being predicted a positive sample is employed since the limited computational accuracy. The highest PCC of 7.31e-01 is given by hybridDBRpred on the DNA-129_Test, which is still quite a difference. The *p*-values against most methods are statistically significant, except that against ESM-NBR and CLAPE on DNA-129_Test are relatively high, which are 3.39e-02 and 7.85e-01, respectively. There are two main potential reasons. First, the identical DNA-binding protein training set; second, both are PLM-based methods like GeSite. The fundamental PLM of ESM-NBR, i.e., ESM2, is trained on UniRef50, while the fundamental PLM of CLAPE i.e., protTrans, is trained on UniRef100. The discriminative features of PLM largely rely on the pretraining dataset. Similar pretraining datasets and supervised learning training sets lead to similar prediction behaviors for native DBS. Nevertheless, the AUC values show that GeSite is much higher than the two control methods in terms of comprehensive prediction performance.

**Table 3.**
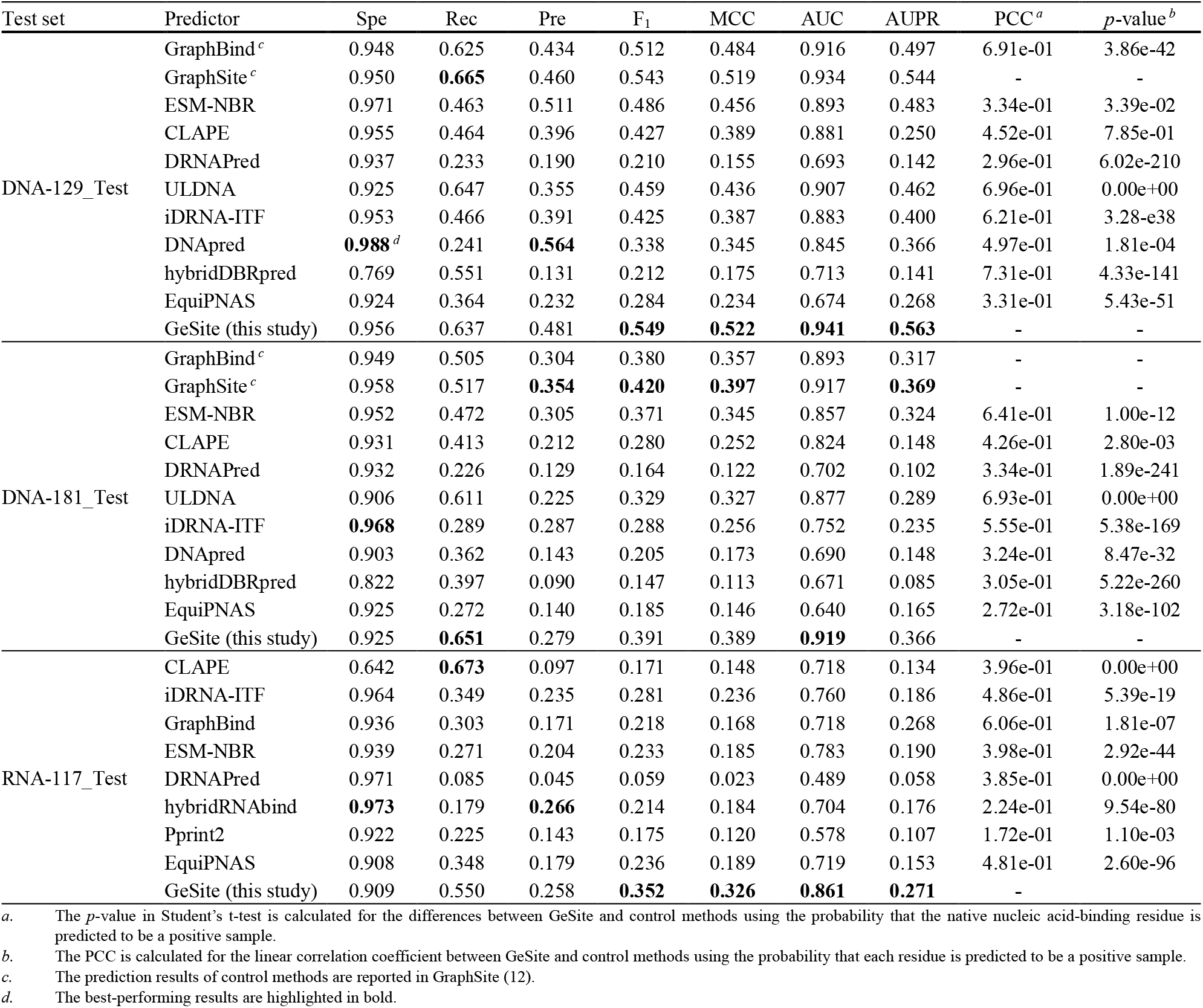
Performance comparison of GeSite and the SOTA NBS prediction methods on three independent test sets.

Apart from that, to show the performance on single residue protein clearly, we plotted scatter plots of head-to-head comparisons of MCC values at single protein (refer to Figure 4). By visiting Figure 4, in most comparisons, GeSite performs better on the vast majority of proteins regardless of the test set. For DBS prediction, taking the result on DNA-181_Test as an example, out of 181 DBPs, there are 134 (74.03%), 132 (72.93%), 105 (58.01%), 169 (93.37%), 125 (69.06%), and 90 (49.72%) cases where GeSite possesses a higher MCC value than six control methods, respectively. For RBS prediction, out of 117 RBPs, there are 58 (49.57%), 90 (76.92%), 82 (70.08%), 96 (82.05%), 90 (76.92%), and 101 (86.32%) cases where GeSite outperforms other predictors separately. These results highlight the excellent performance of the proposed method at the single protein-level. A detailed description of the sources of these prediction results of control methods can be found in Supplementary Text S3.

**Figure 4.**
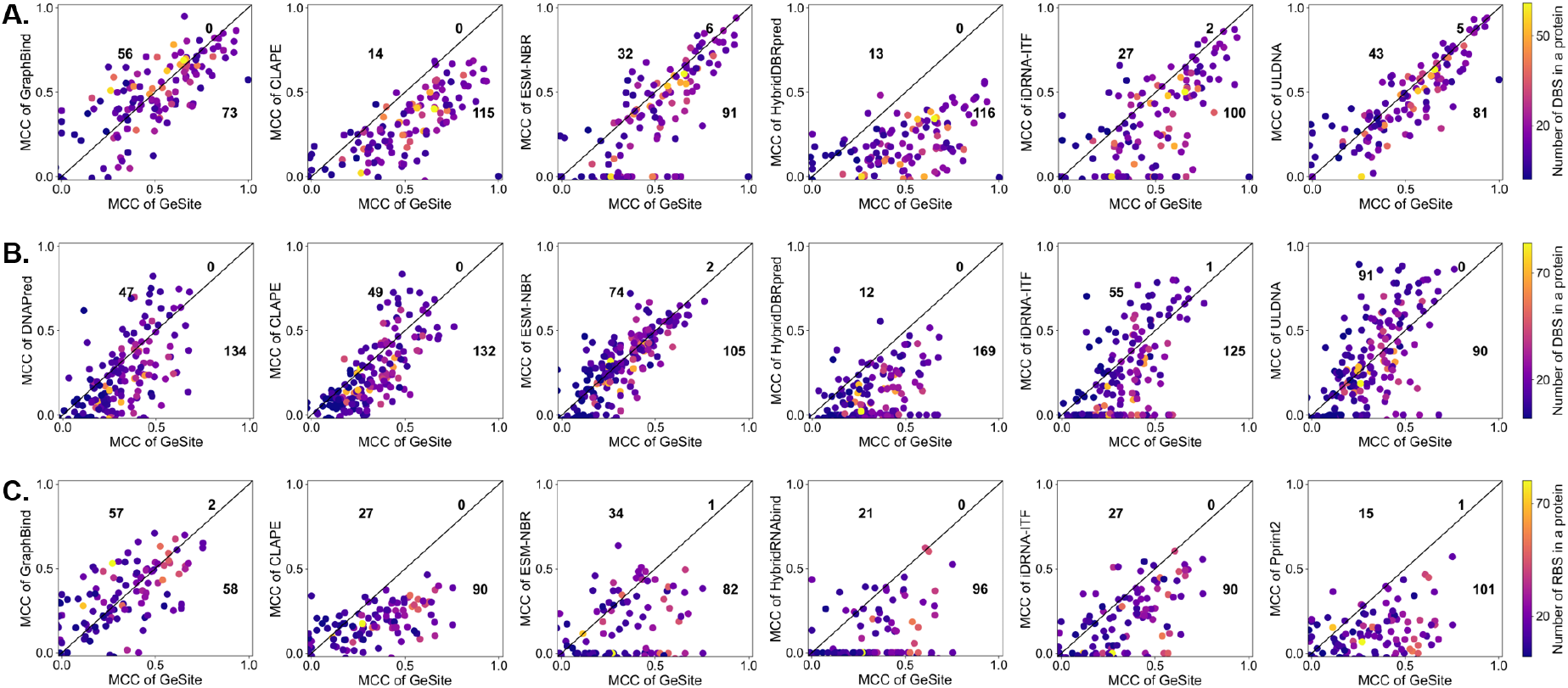
Head-to-head comparison of MCC among GeSite and SOTA methods on three independent test sets. Each point means a test protein. The numbers indicate the number of proteins located in the upper triangle, lower triangle, and diagonal. **(A)**. DNA-129_Test, **(B)**. DNA-181_Test, **(C)**. RNA-117_Test.

### Performance on structurally non-similar proteins

In previous studies, sequences in the test sets similar to those in the training sets were removed using the CH-HIT tool with a clustering threshold of 0.3. While this approach eliminates some structural redundancy, it does not ensure a rigorous one-to-one structural alignment. Given that GeSite and several related works are based on 3D structures, we adopted a more stringent data filtering procedure. Specifically, we employed the US-align tool to calculate the structural identity of each test protein against the training set. Proteins with a maximum TM-score greater than 0.5 were excluded, leaving 49 proteins in the DNA-129_Test set, 92 in the DNA-181_Test set, and 66 in the RNA-117_Test set (see Table S2). Additionally, RNA-285_Test is also included for a more rigorous evaluation.

In Figures 5A-B, GeSite demonstrates the highest MCC values across all four test sets. With the exception of RNA-285_Test, where its performance is comparable to GraphBind, GeSite achieves the highest AUC values for the remaining three test sets. These results underscore the superior generalization capability of our method, particularly on proteins with low structural similarity and newly released proteins.

**Figure 5.**
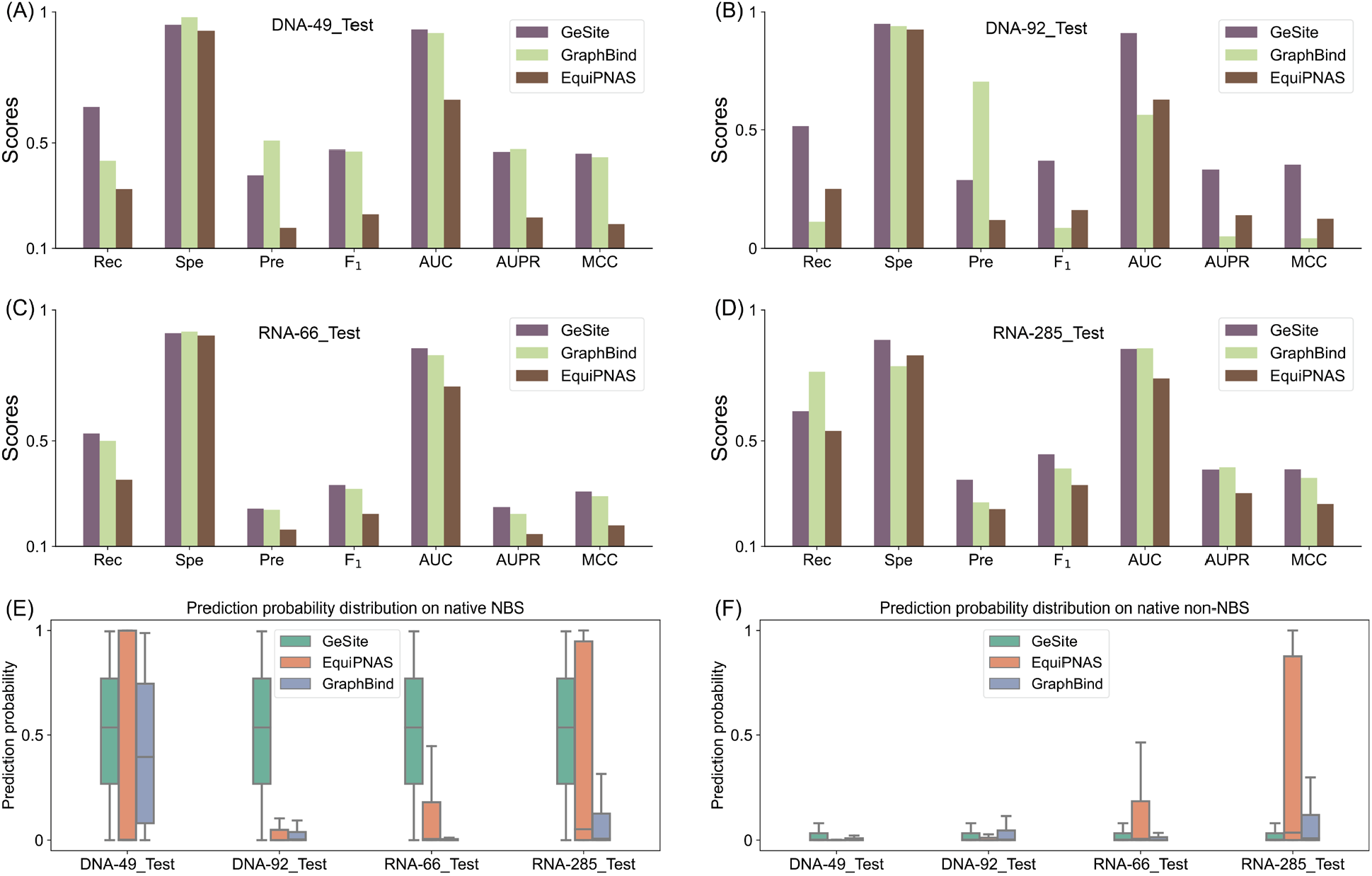
Performance comparison of three SOTA structure-based methods on test proteins with maximum TM-score<0.5 against training data sets. **(A-D)**. evaluation metrics on four test sets; **(E-F)**. prediction probability distributions over positive and negative samples. The vertical axis represents the probability of being predicted as a positive sample. DNA-49_Test, DNA-92_Test, and RNA-66_Test are constructed from the three existing test sets by removing proteins similar to the training set in structure. RNA-285_Test is newly collected in this study from recently released protein-RNA complex.

The predicted probability distribution on the NBS for GeSite is concentrated in the middle, indicating its robustness. In contrast, the distribution for EquiPNAS is broader, which complicates the differentiation between positive and negative samples. GraphBind, on the other hand, exhibits a distribution skewed towards the lower end, favoring negative samples. For negative samples, GeSite maintains a low latitude across all four test sets. While EquiPNAS and GraphBind show similar distributions for both negative and positive samples in RNA-285_Test, this overlap increases the likelihood of confusion between the two classes. These findings further demonstrate the robustness of GeSite in distinguishing between positive and negative samples.

### Case Study

The DNA-binding chain B of human transcription initiation factor TFIID (PDB ID: 6mzm_B, refer to Figure 6A) and the RNA-binding chain A of E3 ubiquitin-protein ligase TRIM71 of Danio rerio (PDB ID: 6fq3_A, refer to Figure 6B) are employed as cases studies. Considering that most of the existing methods need to rely on sufficient MSA information, for a fair assessment, we calculated the MSA depth for both samples using the HHblits tool against the Uniclust30 database with an e-value of 1e-06. We found 459 and 633 homologous sequences for these two proteins, respectively, which are friendly to other predictors.

**Figure 6.**
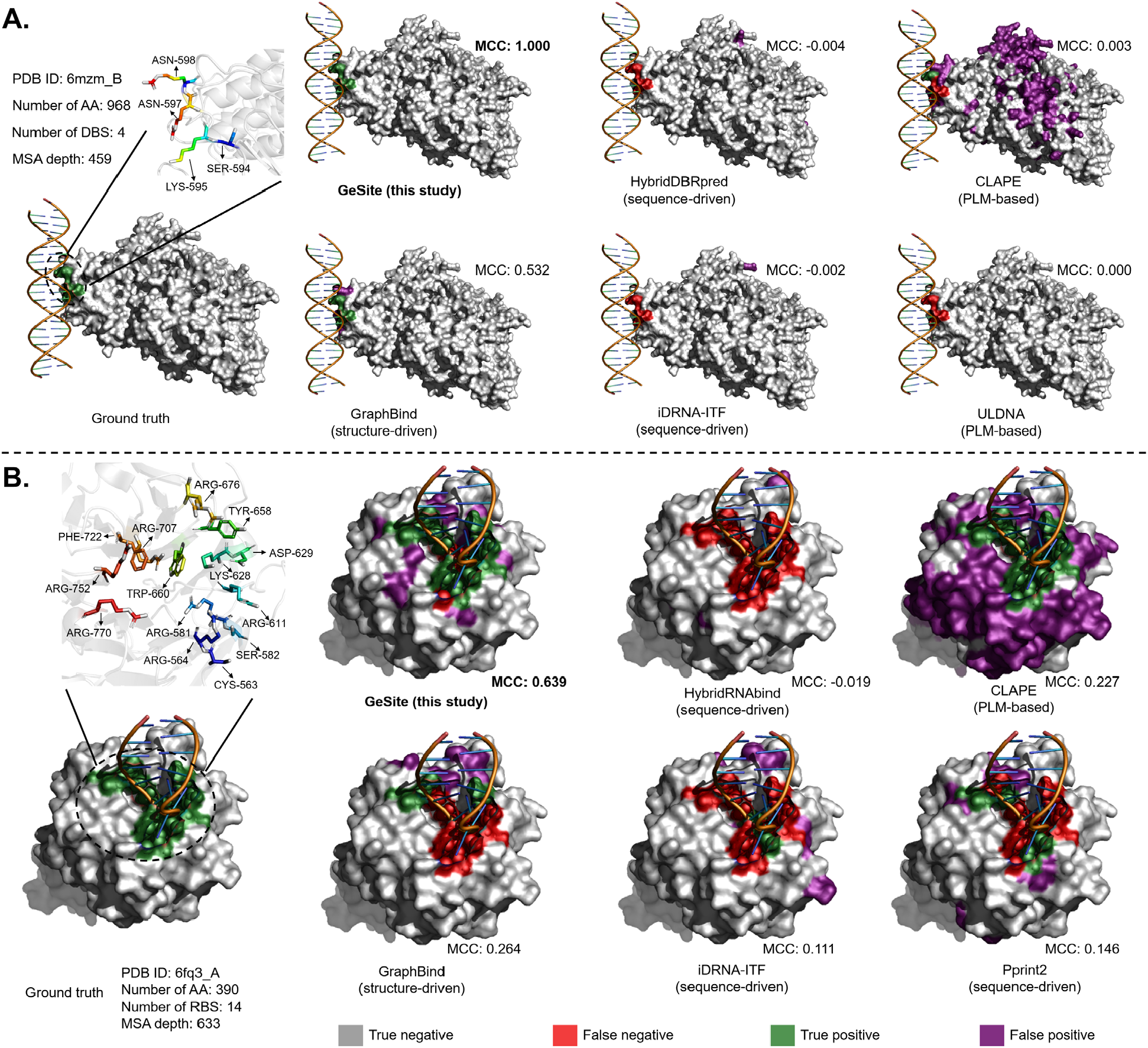
Cases studies on the DNA-binding chain B of human transcription initiation factor TFIID (PDB ID: 6mzm_B) and the RNA-binding chain A of Danio rerio E3 ubiquitin-protein ligase TRIM71 (PDB ID: 6fq3_A). Residues labelled in the upper left corner of each subfigure indicate the native nucleic acid-binding residues. MSA depth is counted by HHblits tool to search Uniclust30 database with an e-value threshold of 1e-06. **(A)**. on 6mzm_B; **(B)**. one 6fq3_A.

From Figure 6A, the prediction of GeSite on 6mzm_B achieves full coverage of the four DBSs, i.e., SER-594, LYS-595, and ASN-597, and ASN-598, while avoiding any incorrect extrapolation of the non-DBS. The MCC value of GeSite is 1.000, which is far beyond any other predictors and 87.96% higher than that of the second-best method, i.e., GraphBind. The four remaining control methods have MCC values around 0. The potential reason for this may be that the sequence of the 6mzm_B is so long (968 amino acids) that it is difficult for these methods to capture discriminating knowledge of the local motif that determines DNA binding although the evolutionary information is sufficient. The advantage of GraphBind over these methods mainly comes from the fact that it is a protein 3D structure-driven method, while other methods rely on MSA information or sequence embedding of PLM from primary sequence only. Tertiary structures that directly guide biological function provide broader discriminatory knowledge for nucleic acid-binding residue prediction. Nevertheless, GraphBind is still confused about the neighboring residues of those natural DBSs, i.e., it predicts those non-DBSs close to the DBSs as positive samples. Instead, GeSite perfectly circumvents such mistakes. This enhancement of GeSite relative to GraphBind may come from augmented pretraining of generic PLM on vast DBP sequences, enabling GeSite to perceive subtle differences of neighboring residues.

In Figure 6B, 6fq3_A presents 14 RBSs, namely, CYS-563, ARG-564, ARG-581, SER-582, ARG-611, LYS-628, ASP-629, TYR-658, TRP-660, ARG-676, ARG-707, PHE-722, ARG-752, and ARG-770, most of which are captured by the proposed GeSite. A MCC value of 0.639 demonstrates the excellence performance of GeSite, which is 0.658, 0.412, 0.375, 0.528, and 0.493 higher than all the control methods respectively. Similar to the DBS prediction, GraphBind also obtains the second-best prediction results, emphasizing the importance of structure information in NBS prediction again. As the sequence-based method, the MCC of PLM-based CLAPE is higher than that of the other two evolutionary information features (typically PSSM and HMM) based-methods, i.e., iDRNA-ITF and Pprint2. This may indicate that PLM provides more useful knowledge for prediction model to identify RBS. The proposed GeSite, which uses 3D structure information and NBS-adaptive PLM, combines the advantages of both to make superior identifications and outperforms other predictors.

### Interpretability and visualization

The ability of proteins to bind nucleic acids derives primarily from a small conserved nucleic acid-binding domain (NBD) capable of highly specific recognition of nucleic acid sequences (1). The GeSite backbone neural network, EGNN, can learn valid knowledge related to spatially neighboring nodes of a single node. Theoretically, the model is able to perceive the specific knowledge of the diverse NBDs from our well-designed protein graph, thus enabling the accurate recognition of NBS. Based on this conjecture, a widely used interpretability analysis algorithm specifically for graph neural networks, i.e. GNNExplainer (47), is employed. For each sample, the output of GNNExplainer contains a protein subgraph, where each node represents those residues that contribute significantly to the prediction of the target residue. Concretely, we selected three DBSs, namely GLN-40 of 5trd_A, ARG-394 of 5ui5_V, and LYS-106 of 6imj_A, as well as two RBSs, namely LYS-140 of 5wty_B, and LYS-271 of 5wwr_A as representatives to find out the key residue nodes for the prediction of these five NBSs.

The analysis results for these three DBSs and two RBSs are shown in Figures 7A-B separately. Apparently, those regions that are significant for identification cover the native NBD. For DBS task, GeSite successfully identifies the DBS GLN-40 of 5trd_A (Thermoplasma acidophilum Riboflavin kinase) as a positive with a score of 0.951. The GNNExplainer algorithm reports 106 salient residues (marked in purple) close to the DNA chain that cover a typical DNA-binding domain (DBD) named Orp-like helix-turn-helix (HTH) domain located at position 13-73 (coated with protein surface). For the residue ARG-394 of 5ui5_V (RNA polymerase sigma factor RpoN of Aquifex aeolicus), GeSite successfully identified it with a probability of 0.843. Similarly, the saliency region of this site, consisting of 150 residues, is highly overlapping with a Homeodomain-like region located at position 376-341. The HTH domain is widely present in a variety of prokaryotic and eukaryotic organisms and plays a fundamental regulatory role (48). The Homeodomain is also known as a classic DBD in eukaryotes. These two cases exemplify the effective utilization of GeSite of near single structure domain discriminative information. In the chain A of 6imj (DNA ligase of African swine fever virus), GeSite presents the attention for multi-domains, both remote and close. Concretely, in the third subfigure of Figure 7A, 6imj_A contains three functional domains, namely N-terminal domain, Active Directory domain, and OB-fold domain, whose positive effects on DNA-protein interactions have been demonstrated in previous study (49). Clearly, the significant residues for prediction of LYS-106 are distributed in all three domains even though the OB-fold domain is far from LYS-106. The ability comes primarily from the ability of the message-passing mechanism of EGNN to allow the network to capture features from remote nodes. These findings further support the idea that GeSite learns valid knowledge from different functional structure domains.

**Figure 7.**
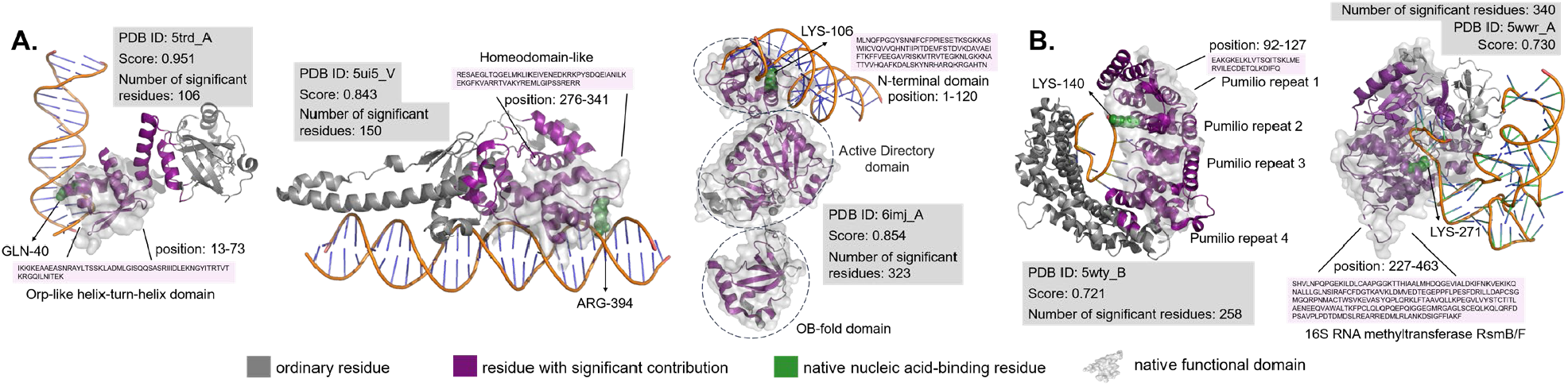
GeSite is enlightened by native nucleic acid-binding domains. Residues selected by GNNExplainer that contribute significantly to the prediction of target NBS (green spheres) are highlighted in purple. The region wrapped by the protein surface is the recorded nucleic acid-binding and related functional domain. The annotations for these domains come from the PDB database (3), which integrates information from several databases such as UniProt, CATH (45), and Pfam (46). Score indicates the probability given by GeSite that the target residue is predicted to be an NBS. **(A)**. on three DBS cases, i.e., GLN-40 in chain A of 5trd, ARG-394 in chain V of 5ui5, and LYS-106 in chain A of 6imj; **(B)**. on two RBS cases, i.e., LYS-140 in chain B of 5wty and LYS-271 in chain A of 5wwr.

On RBS prediction task, similar phenomena are observed. For instance, a prediction score of 0.721 on residue LYS-140 of 5wty_B (Nucleolar protein 9 of Saccharomyces cerevisiae) indicates that GeSite correctly predicted it as an RBS. This chain contains multiple pumilio repeat regions that regulate gene expression by specifically recognizing and binding to RNA sequences (50). The 258 residues with significant contribution for the recognition of LYS-140 cover the pumilio repeat 1 to 4. Likewise, for LYS-271 of chain A of 5wwr (human NSUN6), the purple prominent structure overlaps with 16S RNA methyltransferase RsmB/F region. These findings suggest that, like DBS prediction, GeSite performs identification through the discriminatory information of the RNA-binding domain in the spatial context of the target site. Overall, the above study conveys the idea that the proposed GeSite model is inspired by the NBD in the periphery of the target NBS to be predicted to acquire discriminative knowledge and thus perceive the pattern of nucleic acid-binding. This capability can be extended to multiple functional domains in remote locations.

## CONCLUSIONS

Here, we propose GeSite based on nucleic acid-binding protein domain-adaptive protein language model and E(3)-equivariant graph convolution neural network for accurately predicting nucleic acid-binding residue. Predicted results on multiple independent test sets demonstrate the excellent performance of GeSite. Meanwhile, interpretability analysis on graph neural networks uncovers that the prediction model captures key information about nucleic acid-binding domains thereby helping to identify native nucleic acid-binding residue. While room for improvement remains, GeSite should be an excellent nucleic acid-binding residue prediction tool.

In future research, potential directions of advancement may be the following points: (1). developing a multimodal language model of nucleic acids and proteins to accurately identify proteins-nucleic acid interactions; (2). integrating protein sequence, structure, and function into a single pre-training model to provide higher quality characterization for predictions of downstream tasks; (3). designing a powerful few-shot learning algorithm to precisely predict protein-nucleic acid complex structure directly.

## DATA AVAILABILITY

The wights of GeSite models and RNA-binding protein sequences pretraining data set UniRBP40 are available for academic use at https://huggingface.co/zengwenwu/GeSite/tree/main. The source code, standalone program, DNA- and RNA-binding residues datasets are freely accessible at https://github.com/pengsl-lab/GeSite.

## AUTHOR CONTRIBUTIONS

Wenwu Zeng: Conceptualization, Methodology, Software, Visualization, Writing original draft. Lianrui Pan: Software, Methodology. Boya Ji: Software, Visualization. Liwen Xu: Conceptualization, Methodology, Software, Visualization, Writing original draft. Shaoliang Peng: Funding acquisition, Resources, Writing-review & editing, Supervision.

