## Supplementary Text for "Accurate nucleic acid-binding residue identification based on domain-adaptive protein language model and explainable geometric deep learning"

**Text S1. Evaluation metrics**

We use seven widely used evaluation metrics for assessing binary classification performance, namely, Recall (Rec), Specificity (Spe), Precision (Pre), Matthew's correlation coefficient (MCC), $F_1$-score ($F_1$), the area under operating characteristic curve (AUC), and the area under precision-recall curve (AP). The detailed formulas are shown below:

$$Rec = \frac{TP}{TP+FN} \times 100 \quad (1)$$

$$Spe = \frac{TN}{TN+FP} \times 100 \quad (2)$$

$$Pre = \frac{TP}{TP+FP} \times 100 \quad (3)$$

$$MCC = \frac{TP \cdot TN - FP \cdot FN}{\sqrt{(TP+FN) \cdot (TP+FP) \cdot (TN+FN) \cdot (TN+FP)}} \quad (4)$$

$$F_1 = \frac{2 \cdot TP}{2 \cdot TP + FN + FP} \quad (5)$$

where true positive (TP) and false positive (FP) are the numbers of proteins that are correctly and falsely predicted as nucleic acid-binding residues, and true negative (TN) and false negative (FN) are the numbers of proteins that are correctly and falsely predicted as non-nucleic acid-binding residues respectively. Additionally, Pearson correlation coefficient and *p*-value in Student's t-test are used to calculated the distinctiveness of the proposed method from the previous methods.

**Text S2. ESM-MSA feature extraction**

Unlike normal single-sequence-based protein language model, The Multiple Sequence Alignment (MSA) transformer ESM-MSA (1) built on MSA information digs deeply into the evolutionary relationships within protein families. In this study, the ESM-MSA (1) is employed to generate feature embedding matrices representing the evolutionary information of protein sequences. Specifically, a protein sequence of length $L$ is fed into HHblits tool to generate the raw MSA profile against Uniclust30 database (2) with an e-value of 0.1. Then, we followed the tutorial in https://colab.research.google.com/github/facebookresearch/esm/blob/master/examples/contact_prediction.ipynb and inputted the MSA file into the ESM-MSA model, resulting a feature matrix of size $L \times 768$, where $L$ is the number of residues in protein sequence, and 768 is the feature dimension of each residue.

**Text S3. Sources of predicted results for control methods**

Here we describe the sources of the prediction results of the control methods used for performance comparison with GeSite in detailed. The evaluation metrics of GraphBind (3) and GraphSite were taken directly from the paper of GraphSite (4). The full prediction result of GraphBind for PCC and *p*-value calculation was generated by the standalone package at http://www.csbio.sjtu.edu.cn/bioinf/GraphBind/sourcecode.html. The prediction results of CLAPE (5), ESM-NBR (6), ULDNA (7), Pprint2 (8), and EquiPNAS are generated using the standalone package at https://github.com/YAndrewL/clape, https://github.com/pengsl-lab/ESM-NBR, https://github.com/yiheng-zhu/ULDNA, https://github.com/raghavagps/pprint2, and https://github.com/Bhattacharya-Lab/EquiPNAS, respectively. The prediction results of DRNAPred (9), iDRNA-ITF (10), hybridRNAbind (11), and hybridDBRpred (12) comes from their web servers at http://biomine.cs.vcu.edu/servers/DRNApred/, http://bliulab.net/iDRNA-ITF/, https://www.csuligroup.com/hybridRNAbind/, and http://biomine.cs.vcu.edu/servers/hybridDBRpred/, respectively. The prediction results of DNAPred (13) are contained in result file of hybridDBRpred.

**Table S1.** Performance comparisons of different algorithms on DNA-573_Train and RNA-495_Train over 10-fold cross validation.

| Dataset | Algorithm | | Spe | Rec | Pre | $F_1$ |
|---|---|---|---|---|---|---|
| DNA-573_Train | sequence-based | BiLSTM | 93.53 | 63.69 | 49.51 | 0.557 |
| | | BiGRU | 96.31 | 52.55 | 58.68 | 0.554 |
| | | Self-attention | 91.37 | 60.21 | 41.01 | 0.488 |
| | structure-based | GraphSAGE | 93.59 | 63.22 | 49.56 | 0.555 |
| | | GVP-GNN | 95.40 | 55.65 | 54.66 | 0.551 |
| | | GCN | 81.59 | 65.28 | 26.09 | 0.373 |
| | | GAT | 86.88 | 57.20 | 30.27 | 0.396 |
| | | EGNN | 92.07 | 69.05 | 46.45 | 0.555 |
| RNA-495_Train | sequence-based | BiLSTM | 91.25 | 59.06 | 44.66 | 0.508 |
| | | BiGRU | 91.85 | 52.98 | 43.72 | 0.479 |
| | | Self-attention | 87.64 | 48.93 | 32.12 | 0.387 |
| | structure-based | GraphSAGE | 92.33 | 57.02 | 47.05 | 0.515 |
| | | GVP-GNN | 92.71 | 54.67 | 47.27 | 0.507 |
| | | GCN | 93.76 | 35.73 | 40.63 | 0.380 |
| | | GAT | 87.66 | 49.68 | 32.47 | 0.393 |
| | | EGNN | 92.81 | 55.51 | 48.00 | 0.515 |
